# HOPS-dependent lysosomal fusion controls Rab19 availability for ciliogenesis in polarized epithelial cells

**DOI:** 10.1101/2023.02.07.527563

**Authors:** Huxley K. Hoffman, Rytis Prekeris

## Abstract

Primary cilia are sensory cellular organelles crucial for organ development and homeostasis. Ciliogenesis in polarized epithelial cells requires Rab19-mediated clearing of apical cortical actin to allow the cilium to grow from the apically-docked basal body into the extracellular space. Loss of the lysosomal membrane-tethering HOPS complex disrupts this actin-clearing and ciliogenesis, but it remains unclear how ciliary function of HOPS relates to its canonical function in regulating late endosome-lysosome fusion. Here, we show that disruption of HOPS-dependent lysosomal fusion indirectly impairs actin-clearing and ciliogenesis by disrupting the targeting of Rab19 to the basal body. We also find that Rab19 functions in endolysosomal cargo trafficking apart from its previously-identified role in ciliogenesis. In summary, we show that inhibition of lysosomal fusion abnormally accumulates Rab19 on late endosomes, thus depleting Rab19 from the basal body and thereby disrupting Rab19-mediated actin-clearing and ciliogenesis.

**Summary statement:** Loss of HOPS-mediated lysosomal fusion indirectly blocks apical actin clearing and ciliogenesis in polarized epithelia by trapping Rab19 on late endosomes and depleting Rab19 from the basal body.

## INTRODUCTION

The primary cilium is a sensory organelle, found on most types of cells in vertebrates, which is required for many of the key cellular signaling pathways that coordinate organ development and homeostasis (Anvarian et al., 2019; Goetz and Anderson, 2010; Wheway et al., 2018). Mutations in genes involved in primary cilia formation and function produce genetic disorders termed ciliopathies, affecting multiple organ systems (Hildebrandt et al., 2011; Mitchison and Valente, 2017; Reiter and Leroux, 2017). Renal disease is a prominent feature of many ciliopathies, as defects in the primary cilia of the polarized epithelial cells that line the renal tubules lead to development of kidney cysts and loss of kidney function (Devlin and Sayer, 2019; McConnachie et al., 2021). For example, autosomal dominant polycystic kidney disease, the most common inherited cause of kidney failure, is caused by loss-of-function mutations in a ciliary receptor complex (McConnachie et al., 2021). This makes it important to understand the mechanisms underlying cilia formation in polarized epithelial cells.

Primary ciliogenesis involves extension of a microtubule-based axoneme from a basal body which develops from the mother centriole (Pedersen et al., 2008). The ciliogenesis process shows some mechanistic differences in polarized epithelial cells as compared to other cell types (Hoffman and Prekeris, 2022; Molla-Herman et al., 2010; Sorokin, 1968). In most cell types, the primary cilium initially develops within an intracellular ciliary vesicle which later fuses with the plasma membrane to expose the tip of the cilium to the extracellular space, but in polarized epithelial cells, it appears that the basal body first docks at the apical plasma membrane and then extends the cilium directly into the extracellular space. An important early step in this process is the formation of a clearing in the apical actin cortex (actin clearing) to allow the basal body docking and protrusion and growth of the ciliary axoneme (Hoffman and Prekeris, 2022; Jewett et al., 2021).

Previous work from our lab (Jewett et al., 2021) identified Rab19 GTPase and the homotypic fusion and protein sorting (HOPS) tethering complex as being both required for this apical actin cortical clearing step during ciliogenesis in renal polarized epithelial cells. Rab19 localizes to the site of actin clearing, and knockout (KO) of either Rab19 or the HOPS subunit Vps41 disrupts actin clearing and ciliogenesis. Another HOPS subunit, Vps39, has also been implicated in the regulation of ciliogenesis in renal cells (Iaconis et al., 2020), and all subunits of the HOPS complex were identified in a functional genomic screen for regulators of ciliary signaling (Breslow et al., 2018). The HOPS complex was also shown to be a Rab19 effector, i.e. HOPS interacts with Rab19 in its GTP-bound active state (Jewett et al., 2021).

Rab GTPases in general act as molecular switches that control intracellular membrane trafficking pathways (Homma et al., 2021; Stenmark, 2009), but specific roles of Rab19 apart from its requirement in ciliogenesis remain largely unknown. Among the scarce clues to the overall cellular functions of Rab19 is that the Rab19 ortholog in *Drosophila* localizes to the Golgi (Sinka et al., 2008). The HOPS complex, however, has a well-established role in lysosomal trafficking, where it serves as a membrane tethering complex for fusion of late endosomes (LEs), autophagosomes, and AP-3 vesicles to lysosomes (hereafter referred to collectively as lysosomal fusion) (Balderhaar and Ungermann, 2013; Jiang et al., 2014). Since Rab19 showed little localization to LEs or lysosomes, it was proposed that Rab19 is not involved in lysosomal trafficking and that HOPS interacts with Rab19 to mediate actin clearing and ciliogenesis through a non-lysosomal pathway (Jewett et al., 2021). However, the mechanism by which HOPS regulates ciliogenesis, and whether Rab19 binding to HOPS is directly involved in apical actin clearance, remains unknown.

In this study, we investigated the role of the HOPS complex in actin cortical clearing and primary ciliogenesis. We found that pharmacological inhibition of lysosomal fusion produces similar defects in actin-clearing and ciliogenesis as Vps41 KO, suggesting that the requirement for HOPS in ciliogenesis is a consequence of its requirement in lysosomal trafficking, as opposed to representing a direct non-lysosomal function of HOPS at the basal body. We furthermore found that inhibiting lysosomal fusion impairs ciliogenesis not simply by blocking the degradation of ciliogenesis inhibitors, but rather by disrupting the regulation and localization of key ciliogenesis machinery i.e. Rab19. Additionally, we discovered that Rab19 interacts not only with the late endosome/lysosome (LE/L)-associated HOPS complex but also with the related early endosome (EE)-associated class C core vacuole/endosome tethering (CORVET) complex, and we found that CORVET-containing EEs (in contrast to HOPS-containing LE/Ls) may be directly involved at the basal body in the process of ciliogenesis. Finally, we also found evidence for a previously-uncharacterized role for Rab19 in endolysosomal trafficking. In summary, this study suggests that Rab19 has at least two distinct functions: (1) regulating lysosomal protein traffic; and (2) regulating apical actin clearance and cilia formation in polarized epithelial cells. In contrast, HOPS-depletion appears to affect cilia formation indirectly, predominately by trapping Rab19 on enlarged LEs away from the apically-localized basal body.

## RESULTS

### Disruption of lysosomal fusion impairs apical actin clearing and primary ciliogenesis

The HOPS complex was originally identified as a regulator of lysosomal trafficking, acting as a membrane tether to mediate lysosomal fusion (Balderhaar and Ungermann, 2013; Jiang et al., 2014; Seals et al., 2000; Wartosch et al., 2015). Thus, it might be that HOPS affects ciliation by affecting the lysosomal degradation pathway, or alternatively it might be that HOPS has an additional non-lysosomal function in directly mediating traffic to and from the basal body. Consistent with the former possibility, it has been reported that autophagic degradation of several ciliation inhibitors is needed for primary cilia formation (Liu et al., 2021; Tang et al., 2013; Yamamoto and Mizushima, 2021; Yamamoto et al., 2021). Thus, we first assessed whether the requirement for the HOPS complex in actin cortical clearing and ciliogenesis was related to, or independent of, the role of HOPS in lysosomal fusion. To address this question, we tested whether pharmacological inhibition of lysosomal fusion would produce the same defects in actin clearing and ciliogenesis as Vps41 KO. Chloroquine (CQ) is a lysosomal inhibitor which impairs lysosomal fusion (Mauthe et al., 2018; Mullock et al., 1998), making it a good candidate to pharmacologically mimic the lysosomal fusion defect of Vps41 KO.

We first tested whether CQ treatment had similar effects on the lysosomal pathway as Vps41 KO. We compared the morphology of acidic organelles, i.e. LE/Ls, in Vps41 KO Madin-Darby canine kidney (MDCK) cells to CQ-treated and untreated control wild-type (WT) MDCK cells. LysoTracker dye was used to label all acidic organelles including LE/Ls. Both Vps41 KO and CQ treatment produced a dramatic increase in the size of LysoTracker-positive LE/Ls (Fig. 1A,B). These enlarged LysoTracker compartments likely represent stalled LEs, which have acidified enough for LysoTracker staining, but have failed to fuse to a mature lysosome to degrade their contents and therefore accumulate an abnormal volume of excess cargo. This result supports that CQ treatment has similar effects on the lysosomal pathway as Vps41 KO.

**Figure 1:**
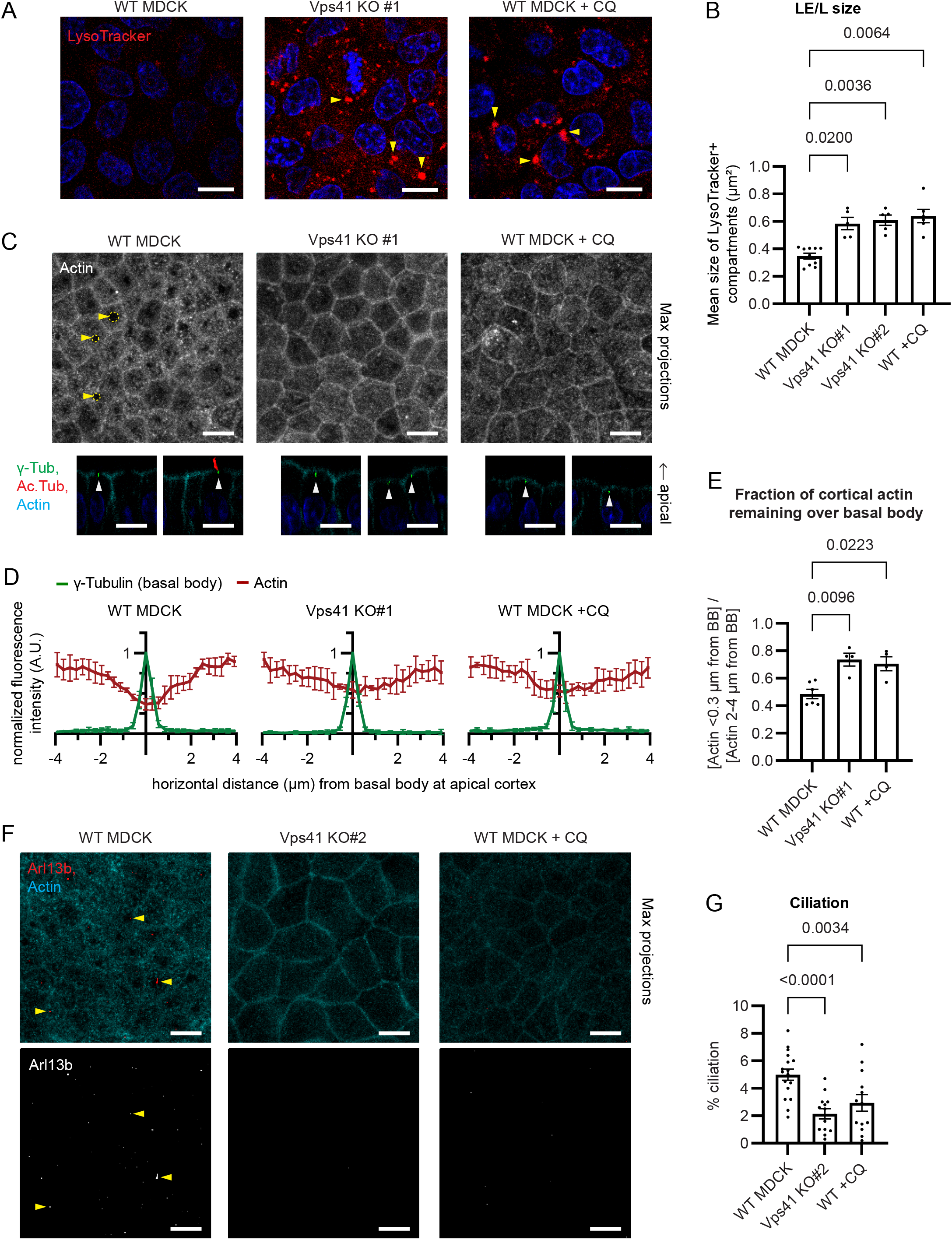
Impaired lysosomal fusion disrupts actin cortical clearing and primary ciliogenesis. A. Parental control wild-type (WT), Vps41 KO, or chloroquine-treated (+CQ) MDCK cells, stained with LysoTracker to label late endosomes and lysosomes (LE/Ls) and Hoechst to label nuclei. Both Vps41 KO and CQ-treated cells show enlarged LE/Ls (examples indicated with arrows). Scale bars 10 μm. Blue color in microscopy images throughout this manuscript shows Hoechst staining of nuclei. B. Mean cross-sectional area of LysoTracker-stained LE/Ls, from experiments as in (A). Each point represents the mean of technical replicates from one biological replicate. Outlier replicates excluded (ROUT = 2%). Error bars: S.E.M. P-values: Brown-Forsythe and Welch ANOVA and Dunnett’s T3 multiple comparisons test. C. WT, Vps41 KO, or CQ-treated MDCK cells, stained with γ-tubulin antibody to label basal bodies, acetylated D-tubulin antibody to label primary cilia, and Phalloidin to label actin. Scale bars 10 μm. Upper panels: maximum intensity projection of Phalloidin; yellow arrows and circles indicate examples of actin cortical clearing. Lower panels: side view slices of merged channels; white arrows indicate basal bodies. Both Vps41 KO and CQ-treated cells show reduced clearing of the apical actin cortex around the basal body. D. Actin intensity profiles, from images as in lower panels of (C). The clearing of the apical actin cortex above the basal body is defective in Vps41 KO and CQ-treated cells. Error bars: S.D. E. Ratio of cortical actin intensity directly above the basal body (within 0.3 μm laterally) to cortical actin intensity flanking the basal body (2-4 μm laterally), from the intensity profiles in (D). Error bars: S.E.M. P-values: Brown-Forsythe and Welch ANOVA and Dunnett’s T3 multiple comparisons test. F. WT, Vps41 KO, or CQ-treated MDCK cells, stained with Arl13b antibody to label primary cilia (arrows indicate examples of cilia) and Phalloidin. Scale bars 10 μm. Both Vps41 KO and CQ conditions show a lack of cilia. G. Percent of cells exhibiting an Arl13b-labeled primary cilium, from experiments as in (F). Both Vps41 KO and CQ treatment impair ciliogenesis. Error bars: S.E.M. P-values: mixed-effects analysis with Geisser-Greenhouse correction and Šidák’s multiple comparisons test.

We then used CQ as a tool to test whether inhibition of lysosomal fusion was sufficient to cause defects in apical actin clearing and ciliogenesis like those observed in Vps41 KO MDCK cells. The reduction in cortical actin above the basal body relative to flanking regions of the apical actin cortex (Fig. 1C-E), and the percent of cells exhibiting a primary cilium after 3 days of culture as confluent monolayers (Fig. 1F,G), were compared between untreated control, CQ-treated, and Vps41 KO cells. Ciliation was assessed after 3 days because cells could withstand this duration of 10 µM CQ without major impacts to cell survival and morphology; the early time point accounts for the low rate of ciliation in WT MDCK as compared to the prior study where ciliation was assessed after 8 days (Jewett et al., 2021). Actin clearing and ciliation were similarly impaired in both Vps41 KO and CQ-treated cells (Fig. 1C-G), indicating that pharmacological inhibition of lysosomal fusion recapitulates the Vps41 KO defects in these processes. These findings suggest that the apical actin-clearing and ciliogenesis defects of Vps41 KO MDCK are likely due to the loss of the canonical HOPS complex function as a membrane tether for lysosomal fusion, rather than reflecting a separate non-lysosomal function of the HOPS complex.

The clearing in the apical actin cortex around the base of the cilium, besides being important to allow for basal body docking and axoneme extension during ciliogenesis (Jewett et al., 2021), has also been shown to regulate the exclusion of non-ciliary apical membrane proteins such gp135 (Francis et al., 2011). Consistent with this, in CQ-treated cells as in Vps41 KO cells where actin clearing was defective, the clearing of gp135 was similarly lost (Fig. 2A). Since both actin clearing and ciliation were reduced but not completely abolished in Vps41 KO and CQ-treated cells, we wondered whether the cilia that did form in these conditions formed within actin clearings, or whether these were cilia aberrantly extended through an intact apical actin cortex. Although the clearing of basal-body-associated cortical actin is impaired in the majority of Vps41 KO and CQ-treated cells, a few cells in these conditions still exhibit normal actin clearing (Fig. 2B), and when cilia were observed in Vps41 KO and CQ-treated cells, they were associated with actin clearings (Fig. 2C). Thus, the occasional cilia found in Vps41 KO or CQ-treated cells represent the occasional cells in which actin clearing was successful, not cells where ciliogenesis proceeded without actin clearing, further demonstrating that ciliogenesis is tightly coupled to actin cortical remodeling. While cilia were typically not found without actin clearings, in contrast, actin cortical clearings without detectable ciliary axonemes were frequently observed (Fig. 1C and Fig. 2D), supporting the model in which actin cortical clearing is required at an early stage of ciliogenesis prior to axoneme extension (Jewett et al., 2021) and thus the actin-clearing defect of Vps41 KO or CQ-treated cells is presumably a cause, as opposed to a consequence, of the ciliation defect.

**Figure 2:**
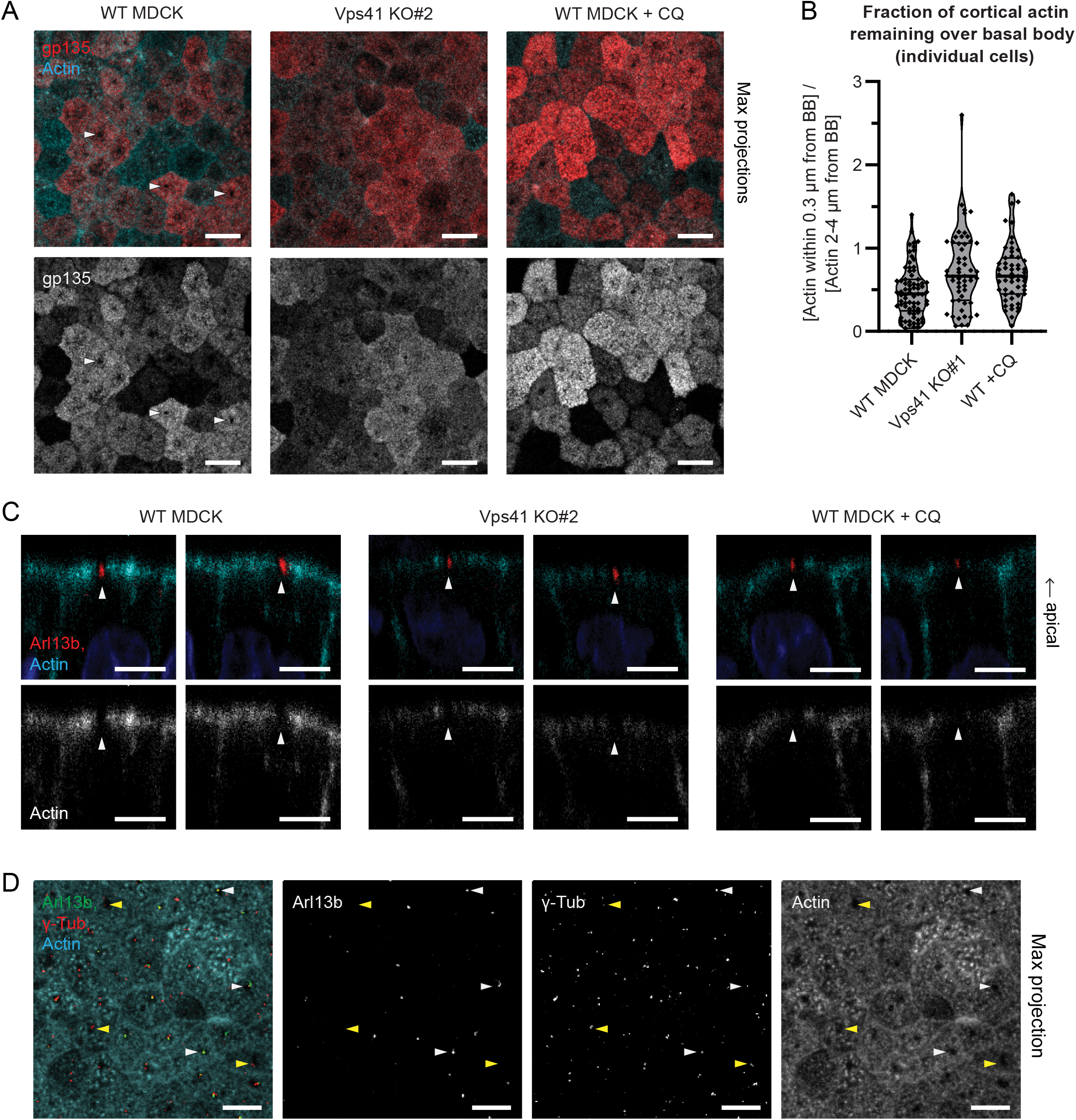
Gp135 cortical clearing and primary ciliogenesis are dependent on actin cortical clearing. A. WT, Vps41 KO, or CQ-treated MDCK cells, stained with gp135 antibody and Phalloidin; maximum intensity projections, scale bars 10 μm. In WT MDCK, the non-ciliary apical membrane protein gp135 is excluded from the site where the apical actin cortex is cleared (examples indicated with arrows). In both Vps41 KO and CQ-treated cells, the lack of a clearing in the actin cortex is accompanied by the lack of an exclusion zone in gp135. B. Fraction of cortical actin remaining above the basal body for all individual cells analyzed, of which the means of each biological replicate are shown in Fig. 1E. Points represent individual cells; bars represent median and quartiles. C. WT, Vps41 KO, or CQ-treated MDCK cells, stained with Arl13b antibody and Phalloidin. Side views, examples selected to show cilia (arrows) although cilia are infrequent overall in Vps41 KO and CQ-treated cells. Scale bars 5 μm. In all three conditions, when a cilium is present, it is associated with a clearing in the apical actin cortex. D. WT MDCK cells stained with Arl13b antibody, γ-tubulin antibody, and Phalloidin; maximum intensity projections, scale bars 10 μm. White arrows indicate examples of cilia in actin clearings, and yellow arrows indicate examples of actin clearings without cilia.

### Ciliation defect of Vps41 KO MDCK is not due to impaired degradation of MYH9, OFD1, or CP110

Previous research has found that autophagy promotes primary ciliogenesis by selectively degrading certain negative regulators of ciliogenesis, including MYH9 (Yamamoto et al., 2021), OFD1 (Tang et al., 2013), and CP110 (Liu et al., 2021). Autophagic degradation relies on HOPS-mediated autophagosome-lysosome fusion (Jiang et al., 2014), and is therefore expected to be impaired by Vps41 KO. We confirmed that Vps41 KO MDCK cells show elevated levels of the autophagosome marker LC3B-II reflecting some defects in autophagic degradation (Fig. S1A,B). We therefore wondered whether the ciliogenesis defects observed in Vps41 KO were due to impaired degradation of MYH9, OFD1, or CP110. MYH9 was reported to inhibit ciliogenesis by stabilizing actin (Rao et al., 2014; Yamamoto et al., 2021), making it particularly likely that a lack of degradation of MYH9 could account for the actin-cortical-clearing defect. We therefore examined MYH9 localization (Fig. S1C) and protein levels (Fig. S1D,E) in Vps41 KO MDCK. If MYH9 were an autophagy cargo and its degradation were blocked by Vps41 KO, then undegraded MYH9 would be expected to accumulate in autophagosomes leading to an overall increase in MYH9 protein levels. We did not observe accumulation of GFP-MYH9 in LC3B-labeled autophagosomes in Vps41 KO MDCK cells or in MDCK cells treated with the autophagy inhibitor Bafilomycin A1 (BafA) (Fig. S1C), nor were MYH9 levels significantly altered in Vps41 KO MDCK (Fig. S1D,E), suggesting that MYH9 does not appear to be an autophagy cargo in MDCK cells under the conditions of these experiments. We also examined the effects of GFP-MYH9 overexpression in WT MDCK cells and found that elevated levels of MYH9 did not block actin clearing or ciliation in these cells (Supp Fig. S1F). Thus, contrary to previous reports in RPE1, IMCD3 cells (Rao et al., 2014), and MEFs (Yamamoto et al., 2021), MYH9 did not appear to be a ciliogenesis inhibitor in MDCK cells. Vps41 KO also showed little effect on cellular OFD1 or CP110 protein levels (Fig. S1D,E). Our data therefore did not support the hypothesis that the actin-clearing and ciliogenesis defects of Vps41 KO cells were due to impaired autophagy of MYH9, OFD1, or CP110.

### Disruption of lysosomal fusion impairs ciliogenesis by retaining Rab19 on enlarged LEs and sequestering it away from the basal body

Since Rab19 interacts with the HOPS complex and functions in actin cortical clearing and ciliogenesis (Jewett et al., 2021), we wondered whether the ciliogenesis defects caused by inhibition of lysosomal fusion were due to an effect on Rab19. We therefore examined whether Rab19 localization was altered in Vps41 KO or CQ-treated cells. As previously reported (Jewett et al., 2021), in WT MDCK, Rab19 vesicles were enriched around the basal body and actin clearing site (Fig. 3A,B), and Rab19 showed little localization to LE/Ls (Fig. 3C,D). We also observed a substantial pool of Rab19 in the Golgi (Fig. S2A,B), consistent with the reported Golgi localization of Rab19 in *Drosophila* (Sinka et al., 2008). Although the Golgi apparatus localizes around the centrosome in many mammalian cell types (Masson and El Ghouzzi, 2022), we noted that in polarized MDCK monolayers the Golgi was not detected at the actin-clearing site around the apically docked basal body, but rather closer to the nucleus (Fig. S2B), so the basal-body localization of Rab19 appeared to be separate from its Golgi localization.

**Figure 3:**
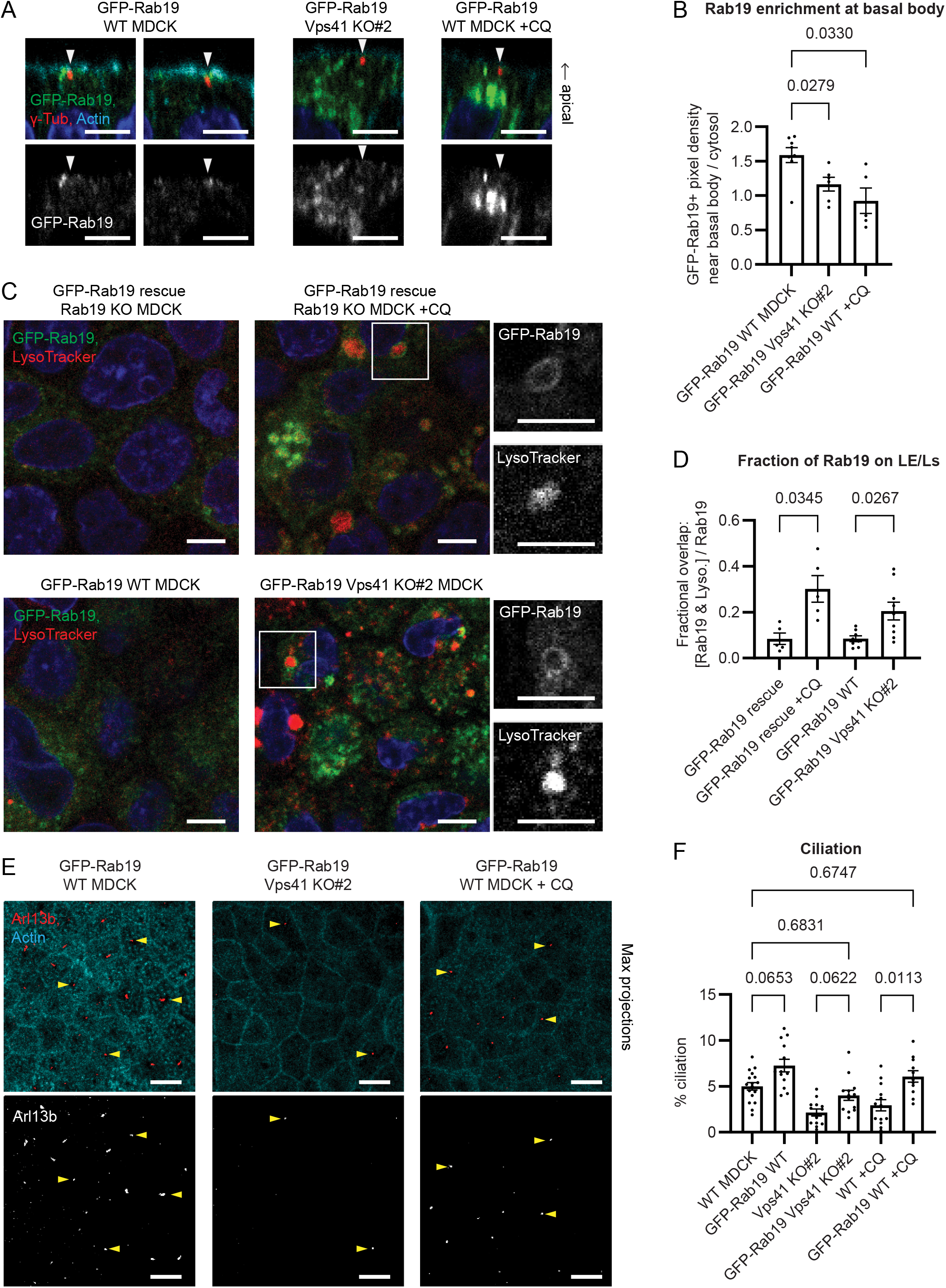
Disruption of lysosomal fusion impairs ciliogenesis by relocalizing Rab19 to enlarged LEs away from the basal body. A. MDCK cells expressing GFP-Rab19 on WT background with or without CQ treatment, or GFP-Rab19 on Vsp41 KO background, stained with γ-tubulin antibody, Phalloidin, and Hoechst. Side views, scale bars 5 µm. Basal-body localization of Rab19 is decreased in Vps41 KO and CQ-treated cells. B. Fold enrichment of GFP-Rab19 at the basal body, from experiments as shown in (A). Error bars: S.E.M. P-values: Brown-Forsythe and Welch ANOVA and Dunnett’s T3 multiple comparisons test. C. MDCK cells expressing GFP-Rab19 on a Rab19 KO background with or without CQ treatment (upper panels), or GFP-Rab19 on a WT or Vps41 KO background (lower panels), stained with LysoTracker. Scale bars 5 µm. Rab19 shows little localization to LE/Ls under basal conditions, but accumulates on membranes of enlarged LE/Ls in CQ-treated or Vps41 KO cells. D. Fractional overlap between GFP-Rab19 and LysoTracker, from experiments as in (C). Error bars: S.E.M. P-values: Brown-Forsythe and Welch ANOVA and Dunnett’s T3 multiple comparisons test. E. GFP-Rab19 overexpressing WT, Vps41 KO and CQ-treated MDCK cells, stained with Arl13b antibody and Phalloidin. Maximum intensity projections, scale bars 10 µm. Arrows point to examples of cilia. Compare to examples of the non-GFP-Rab19-expressing parental cell lines in Fig. 1. F. Percent of cells with an Arl13b-labeled primary cilium, from experiments as in (E). Data for the non-GFP-Rab19-expressing cell lines is the same as in Fig. 1, repeated here for comparison to the GFP-Rab19-overexpressing cell lines. Error bars: S.E.M. P-values: mixed-effects analysis with Geisser-Greenhouse correction and Šidák’s multiple comparisons test. GFP-Rab19 overexpression enhanced ciliation in WT MDCK, partially rescued ciliation in Vps41 KO, and fully rescued ciliation in CQ-treated cells.

In Vps41 KO or CQ-treated MDCK cells, however, Rab19 localized strongly to the membranes of enlarged LE/Ls (Fig. 3C,D). Importantly, this was accompanied by a reduction in basal-body enrichment of Rab19 (Fig. 3A,B). These results indicate that impairment of lysosomal fusion results in aberrant retention of Rab19 on LEs, thus, depleting the pool of Rab19 available to mediate actin clearing and ciliogenesis at the basal body. The portion of Rab19 localizing in the Golgi was also reduced in Vps41 KO or CQ-treated cells, although these conditions did not appear to grossly disrupt the structure of the Golgi itself (Fig. S2A,B), suggesting that the accumulation of Rab19 on LEs sequestered Rab19 away from its normal sites of action at both the basal body and the Golgi.

To assess whether depletion of Rab19 from the basal body was the mechanism for the actin-clearing and ciliogenesis defects in Vps41 KO and CQ-treated cells, we tested whether overexpression of Rab19 could rescue the defects. Indeed, GFP-Rab19 overexpression fully rescued actin clearing and ciliation in CQ-treated cells and partially rescued these phenotypes in Vps41 KO cells (Fig. 3E,F), and overexpression of GFP-Rab19 also enhanced ciliation in control cells (Fig. 3E,F), supporting the hypothesis that mislocalization of Rab19 contributes to the actin-clearing and ciliogenesis defects produced by inhibition of lysosomal fusion (either by Vps41-KO or by CQ treatment).

### Rab19 functions in cargo transport to LEs, and interacts with V-ATPase

The observation that Rab19 accumulates on LEs in Vps41 KO or CQ-treated cells led us to investigate how Rab19 is involved in the endolysosomal pathway. Since Rab19 showed little localization to LE/Ls at steady state but accumulated on LEs when their fusion was blocked (Fig. 3C,D), it is likely that under normal conditions Rab19 is transiently recruited to LE membranes and then released from those membranes following lysosomal fusion. To investigate the function of Rab19 targeting to LE/Ls, we examined the LE/L phenotypes for Rab19 KO and for the Rab19 constitutively active (CA) mutant Q79L in MDCK cells. Neither Rab19 KO nor Rab19-CA showed any major effect on LE/L size under steady-state conditions (Fig. S2C,D). However, Rab19-CA expression greatly exaggerated the enlarged LE/L phenotype induced by CQ (Fig. S2C,D). A likely interpretation for this finding is that Rab19 may mediate trafficking of certain cargoes to LE/Ls. Under conditions of normal lysosomal function, an altered rate of delivery of those cargoes may have limited effect on LE/L size because the cargo is rapidly degraded in lysosomes, and thus it is only when lysosomal degradation is inhibited (e.g. by CQ treatment) that increased delivery of certain cargoes results in increased swelling of the stalled LE/Ls.

Interestingly, the recruitment of Rab19 to LEs did not appear to rely on its binding to the HOPS complex, since it was observed in the Vps41 KO cells (Fig. 3C,D) in which HOPS complex formation is disrupted (Ostrowicz et al., 2010) and the remainder of HOPS does not localize to LEs (Fig. S2E). This suggests that Rab19 is targeted to the LE by some other factor (or alternatively Rab19 could be targeted to the EE and remain associated with the endosome as it matures to LE) and may subsequently interact with the HOPS complex on the LE membrane.

Examining previously published proteomics analysis of Rab19 binding partners (Jewett et al., 2021) for factors relating to the endolysosomal pathway, we noted that in addition to the HOPS complex, Rab19 also interacted with several subunits of the cytosolic V1 sector of the vacuolar-type ATPase (V-ATPase). The V-ATPase is a proton pump which assembles on endolysosomal membranes to acidify these organelles (McGuire et al., 2017; Podinovskaia and Spang, 2018), and has also been shown to act as a pH sensor mediating recruitment of certain cytosolic trafficking factors to endosomal membranes in an manner dependent on the intra-endosomal pH (Hurtado-Lorenzo et al., 2006; Marshansky et al., 2014). In contrast to the GTP-dependent binding of Rab19 to HOPS, the proteomics results indicated that Rab19 binding to the V-ATPase subunit ATP6V1A was GTP-independent (Jewett et al., 2021), suggesting that ATP6V1A can interact with both GTP-bound (active and membrane-associated) and GDP-bound (inactive and cytosolic) forms of Rab19. Using GFP nanobody to pull-down GFP-Rab19 from MDCK cell lysate, we confirmed that Rab19 does interact with ATP6V1A in a GTP-independent manner (Fig. S2F). The V-ATPase could therefore be a plausible candidate to regulate the transient recruitment of Rab19 to LEs.

### Rab19 interacts with core subunits of both HOPS and CORVET complexes

To explore the significance of the Rab19-HOPS interaction, we set out to identify which subunit of the HOPS complex mediates its binding to Rab19. The HOPS complex contains two HOPS-specific subunits (Vps41 and Vps39) along with four core subunits (Vps11, Vps16, Vps18, and Vps33A) which are shared with the early endosomal CORVET complex (Balderhaar and Ungermann, 2013) (Fig. 4A).

**Figure 4:**
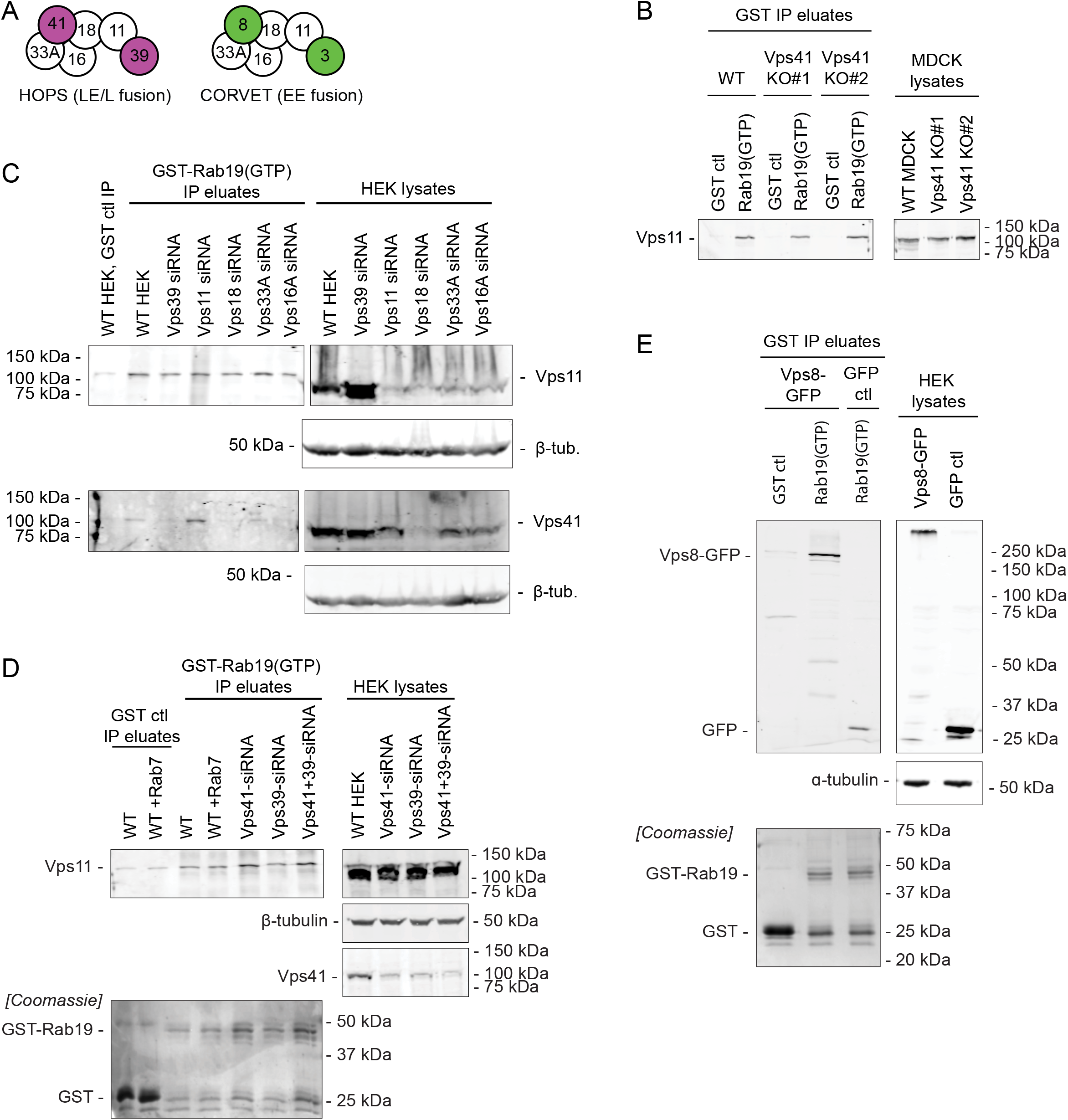
Rab19 interacts with the HOPS/CORVET core complex. A. Vps protein subunits of the HOPS and CORVET complexes. HOPS-specific subunits shown in purple, CORVET-specific subunits in green, and shared core subunits in white. B. Anti-Vps11 western blot of GST-Rab19 immunoprecipitation (IP) assay on WT and Vps41 KO MDCK lysates, showing that Rab19 interacts with Vps11 regardless of presence of Vps41. C. Anti-Vps11 and anti-Vps41 western blots of GST-Rab19 IP assay on WT and Vsp39-, Vps11-, Vps16-, Vps18-, and Vps33A-siRNA HEK293T lysates. Knockdown of any one of the core subunits severely disrupted Vps11 and Vps41 expression levels. D. Anti-Vps11 and anti-Vps41 western blots of GST-Rab19 IP assay on WT and Vsp39-, Vps41-, and Vps39/41-siRNA HEK293T lysates, and on HEK293T lysates with addition of purified recombinant Rab7, showing that Rab19-Vps11 interaction is not dependent on Vps39 or Vps41 and is not competitive with Rab7. E. Anti-GFP western blot of GST-Rab19 IP on HEK293T cells transfected either with Vps8-GFP or with GFP control, showing that Rab19 interacts with CORVET subunit Vps8.

As previously reported (Jewett et al., 2021), glutathione bead pull-down experiments with recombinantly produced GST-Rab19 confirmed that Rab19 in its GTP-bound state interacts with HOPS (as determined by immunoblotting for the Vps11 subunit) in WT MDCK cell lysates (Fig. 4B and Fig. S3A). Importantly, we also detected the interaction of Rab19 with Vps11 in Vps41 KO MDCK lysates (Fig. 4B and Fig. S3A), suggesting that Rab19 binding to HOPS complex is not mediated by Vps41. To probe the dependence of Rab19-HOPS interaction on the other HOPS subunits, we used siRNA to deplete each of the HOPS subunits in HEK293T cells (Fig. 4C,D, and Fig. S3B-E). Knockdown of any one of the four HOPS/CORVET core subunits severely disrupted the entire HOPS complex, as indicated by decreased levels of Vps11 and Vps41 in cell lysates (Fig. 4C and Fig. S3B,F), so the roles of these core subunits in Rab19-HOPS binding could not be individually assessed. Knockdown of Vps39, in contrast, had only moderate effects on Vps11 protein levels, and did not impair the interaction of Rab19 with Vps11 (Fig. 4C,D, and Fig. S3B-D,F). Thus, neither one of the HOPS-specific subunits, Vps41 or Vps39, appear to be required for Rab19-HOPS interaction.

The HOPS complex has been shown to bind Rab7 *via* both Vps41 and Vps39 (Brett et al., 2008; Bröcker et al., 2012). We wondered whether Rab19 might similarly interact with both Vps41 and Vps39, which could explain why knockdown of either Vps41 or Vps39 alone failed to disrupt the interaction. However, co-depletion of both Vps41 and Vps39 still did not reduce Rab19 binding to Vps11 (Fig. 4D and Fig. S3D). We further investigated whether Rab19 competes with Rab7 for binding to HOPS, by testing whether addition of purified recombinant Rab7 in the GST-Rab19 pulldown experiment would reduce Rab19-HOPS binding. Addition of Rab7 did not impair Rab19-HOPS interaction (Fig. 4D and Fig. S3D,G), suggesting that Rab19 binds to HOPS at a different site than Rab7 does. All these results indicate that Rab19-HOPS interaction is independent of HOPS-specific subunits, suggesting that HOPS core subunits may mediate interaction with Rab19.

Since HOPS and CORVET share the same core subunits, we therefore wondered whether Rab19 also interacts with the CORVET complex. To test this possibility, we performed GST-Rab19 pull-down assays using lysates of HEK293T cells expressing CORVET-specific subunit Vps8-GFP. Vps8-GFP (presumably as part of CORVET complex) did bind to GST-Rab19 (Fig. 4E and Fig. S3H), supporting the hypothesis that Rab19 can also interact with CORVET. Collectively, these results suggest that Rab19 likely binds to HOPS/CORVET core subunits, and thus, likely interacts not only with HOPS but also with CORVET complex.

### EEs but not LE/Ls are found at the site of ciliogenesis

If Rab19 binds to both HOPS and CORVET, we wondered whether either HOPS or CORVET are directly involved in the ciliogenesis function of Rab19. To test this question, we examined the subcellular localization of the HOPS/CORVET core by immunostaining for Vps11. Vps11 puncta were frequently observed at the periphery of the actin cortical clearing (Fig. 5A,B,D), where they sometimes overlapped with Rab19 (Fig. 5A), potentially supporting a direct role for HOPS and/or CORVET in ciliogenesis.

**Figure 5:**
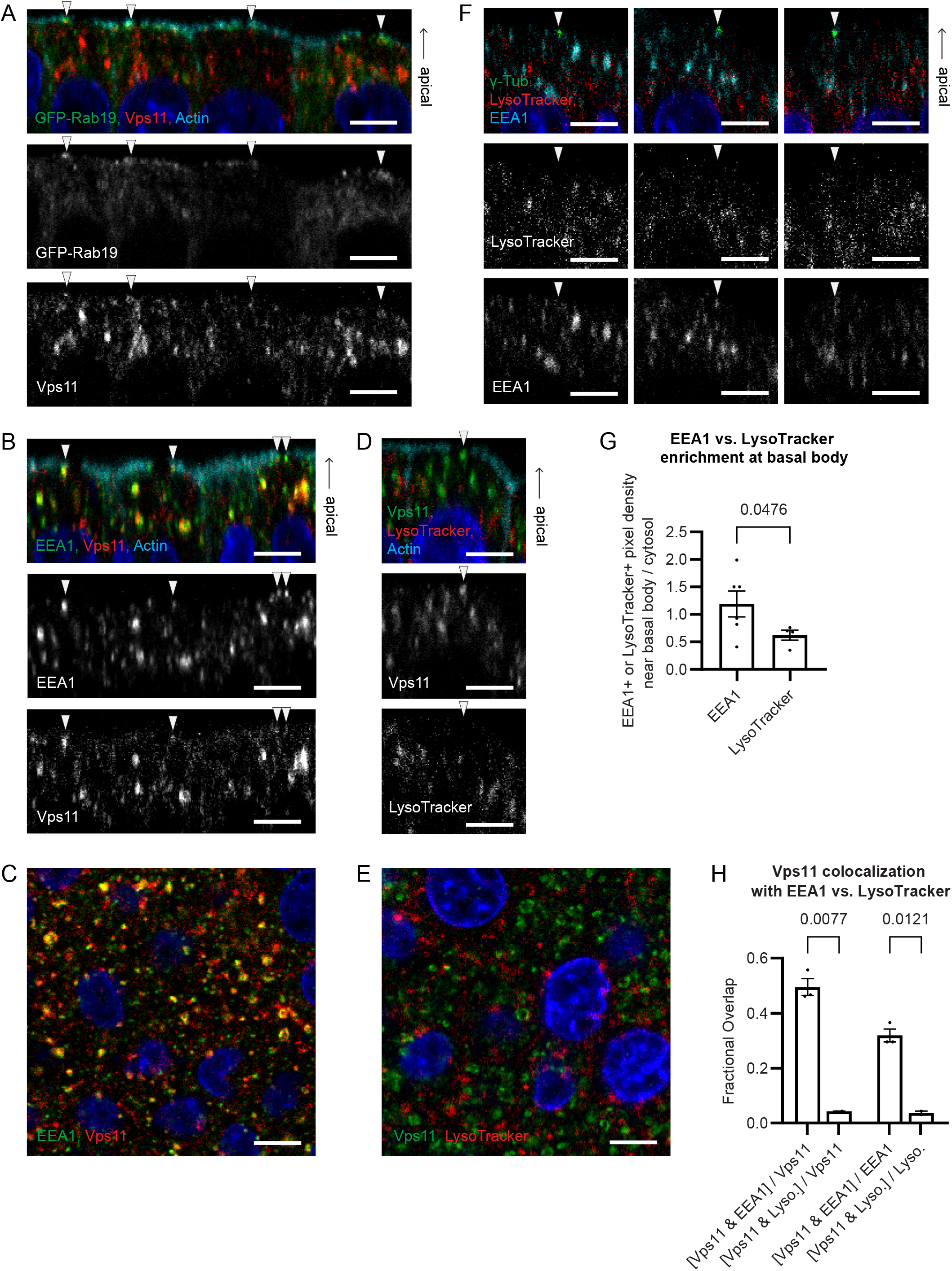
CORVET-containing EEs are found at the site of ciliogenesis. A. GFP-Rab19 overexpressing WT MDCK cells stained with Vps11 antibody and Phalloidin; side view, scale bars 5 µm. Vps11 overlaps with Rab19 at the periphery of the actin clearing (arrows). B. WT MDCK cells stained with Vps11 and EEA1 antibodies and Phalloidin; side view, scale bars 5 µm. Vps11 at the actin clearing is co-labeled with EEA1 (arrows). C. WT MDCK cells stained with Vps11 and EEA1 antibodies. Scale bars 5 µm. Vps11 colocalizes with EEA1. D. WT MDCK cells stained with Vps11 antibody, LysoTracker, and Phalloidin; side view, scale bars 5 µm. Vps11 at the actin clearing (arrow) has little to no co-labeling with LysoTracker. E. WT MDCK cells stained with Vps11 antibody and LysoTracker. Scale bars 5 µm. Vps11 has little colocalization with LysoTracker. F. WT MDCK cells stained with γ-tubulin and EEA1 antibodies and LysoTracker; side view, scale bars 5 µm. EEA1 but not LysoTracker is observed at the basal body. G. Fold enrichment of EEA1 or LysoTracker at the basal body, from experiments as in (F). Error bars: S.E.M. P-value: Kolmogorov-Smirnov test. H. Fractional overlap of Vps11 antibody with EEA1 antibody or with LysoTracker, from images as in (C) and (E). The majority of Vps11 is colocalized with EEA1 and not with LysoTracker. Error bars: S.E.M. P-values: Brown-Forsythe and Welch ANOVA and Dunnett’s T3 multiple comparisons test.

We were unable to directly assess the distinct localizations of HOPS and CORVET, because commercially available antibodies for the HOPS-specific and CORVET-specific subunits did not perform adequately for immunofluorescence in MDCK cells, and overexpression of tagged HOPS subunits disrupts the function of these complexes in ciliogenesis (unlike Rab19, for which overexpression of a tagged form enhances ciliation in WT cells and rescues ciliation in Rab19 KO, indicating that tagged Rab19 is functional in ciliogenesis) (Jewett et al., 2021). To distinguish whether the Vps11 detected at the site of ciliogenesis represented HOPS or CORVET, we therefore used immunostaining for early endosome antigen 1 (EEA1) as a marker for EEs (organelles containing CORVET), and LysoTracker dye as a marker for LE/Ls (organelles containing HOPS). EEA1, like Vps11, was often detected at the basal body and actin cortical clearing (Fig. 5B,F,G), and Vps11 colocalized strongly with EEA1, both at the ciliation site and overall in the cells (Fig. 5B,C,H). In contrast, LysoTracker showed little localization to the site of ciliogenesis (Fig. 5D,F) and little colocalization with Vps11 (Fig. 5D,E,H). Thus, it appears that in WT MDCK the majority of Vps11, including the Vps11 present at the site of ciliogenesis, is part of CORVET complex on EEs, with relatively little Vps11 in HOPS complex on LE/Ls.

To probe whether EEs function in ciliogenesis or in subsequent trafficking to or from cilia, we also examined EEA1 localization with respect to Arl13b. EEA1-positive EEs were frequently found at sites of actin cortical clearing that did not show cilia, and also at the base of Arl13b-positive cilia (Fig. S4A), suggesting that these EEs may be involved both early in ciliogenesis and later in ciliary trafficking. We then wondered whether Rab19 and its interaction with CORVET was responsible for targeting EEs to the ciliogenesis site. Rab19 KO MDCK cells also showed EEA1 at the basal body (Fig. S4B), indicating that Rab19 is not required for localizing EEs to this site. It may instead be that Rab19 transports some ciliogenesis factors to the EEs at the basal body to promote ciliogenesis. We also tested whether EEs at the site of ciliogenesis were perturbed by inhibition of lysosomal fusion. EEA1 was observed at basal bodies in Vps41 KO and CQ-treated as well as WT MDCK (Fig. S4C), indicating that inhibition of lysosomal fusion did not disrupt the targeting of EEs to the site of ciliogenesis, although it might well be that the contents of these EEs are dysregulated by accumulation of cargoes that would normally be transported to lysosomes and degraded.

These observations support the idea that the interaction of CORVET with Rab19 might play a direct role in ciliogenesis, although further study will be needed to demonstrate that role. In contrast, we see little evidence for HOPS or LE/Ls at the site of ciliogenesis (albeit we cannot rule out the possibility that small LE/Ls with LysoTracker signal below the detection limit of this assay could be present there), suggesting that HOPS and its interaction with Rab19 are likely not directly involved in ciliogenesis. This is consistent with the model in which the ciliogenesis defects of Vps41 KO or CQ-treated cells are due to aberrant retention of Rab19 on LEs impeding recruitment of Rab19 to the site of ciliogenesis (Fig. 6).

**Figure 6:**
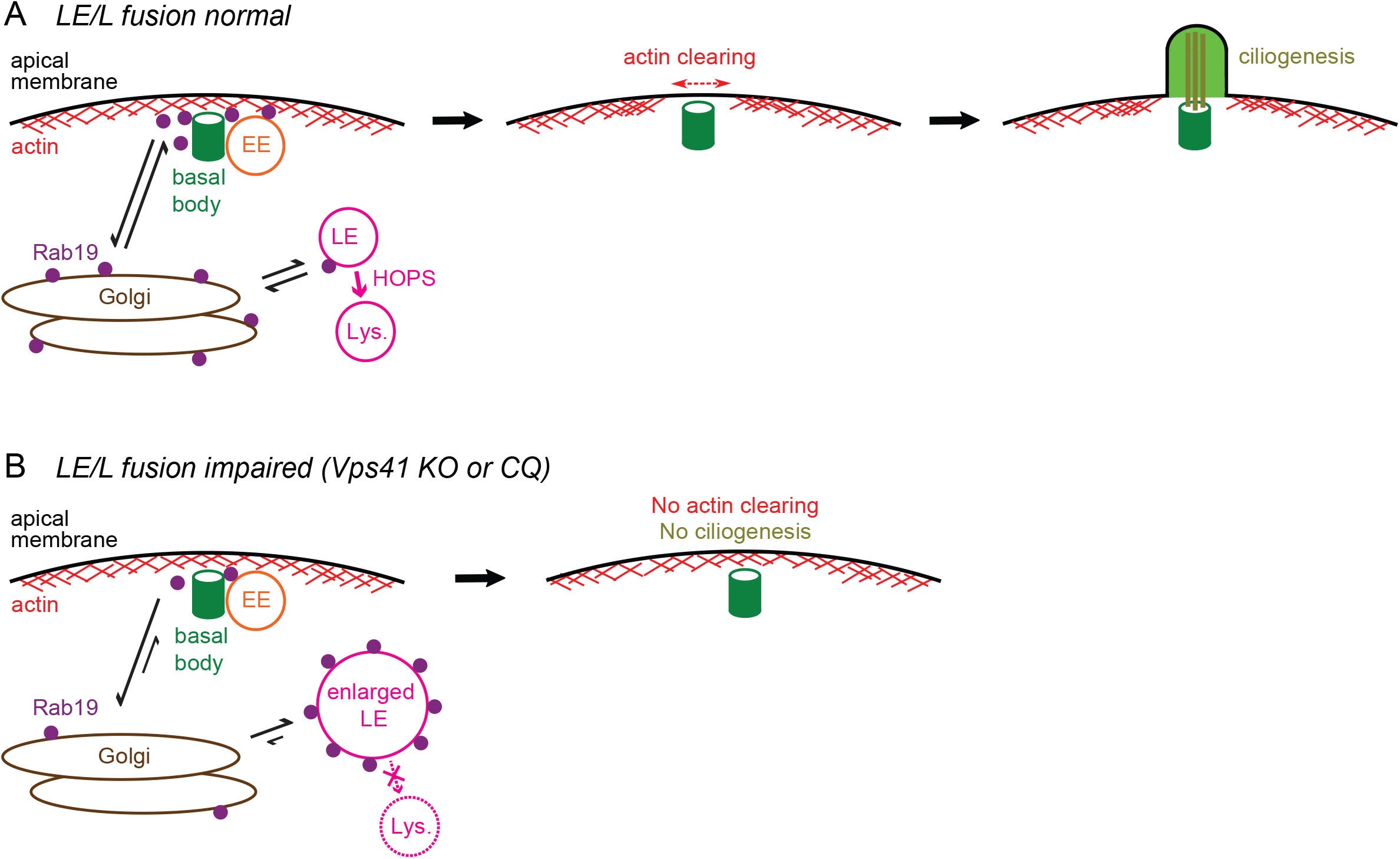
Model: HOPS-dependent lysosomal fusion regulates actin cortical clearing and ciliogenesis by controlling Rab19 availability at the basal body. A. Under normal conditions, a large portion of Rab19 is targeted to the basal body to drive apical actin cortical clearing and ciliogenesis, while a minor population of Rab19 is transiently recruited to LE membranes and then cycles off of these membranes following lysosomal fusion. EEs, but not LE/Ls, are enriched at the basal body. B. When lysosomal fusion is impaired by Vps41 KO or CQ treatment, then Rab19 is not released from LEs, which become enlarged with undegraded cargo. Accumulation of Rab19 on LEs depletes the pool of Rab19 that can be recruited to the basal body, leading to defects in actin-clearing and ciliogenesis.

## DISCUSSION

In this study, we set out to elucidate the role of the HOPS tethering complex in apical actin cortical clearing and primary ciliogenesis. Knockout or mutation of HOPS subunits was recently shown to disrupt primary ciliogenesis and ciliary signaling (Breslow et al., 2018; Iaconis et al., 2020; Jewett et al., 2021). In polarized epithelial MDCK monolayers, Vps41 KO was shown to block a key early step of ciliogenesis in which the apical cortical actin is remodeled and cleared around the apically docked basal body to allow the ciliary axoneme to grow into the extracellular space (Hoffman and Prekeris, 2022; Jewett et al., 2021). The canonical role of the HOPS complex is as a membrane tether that mediates lysosomal fusion and is thus required for lysosomal degradation (Balderhaar and Ungermann, 2013; van der Beek et al., 2019). It was unclear whether the requirement for HOPS in ciliogenesis reflected a requirement for lysosomal trafficking in ciliogenesis, or whether it represented a separate function of the HOPS complex. It was also unknown whether HOPS participates directly in actin-clearing and ciliogenesis at the basal body, or whether its role is indirect. Addressing these outstanding questions is the main focus of this study.

### Requirement for lysosomal trafficking in ciliogenesis

We reasoned that if the actin-clearing and ciliogenesis role of HOPS were a consequence of its role in lysosomal fusion, then treatment with CQ, a pharmacological inhibitor of lysosomal fusion (Mauthe et al., 2018; Mullock et al., 1998), should recapitulate the actin-clearing and ciliogenesis defects of Vps41 KO. CQ had similar effects on LE/Ls in MDCK cells as Vps41 KO, producing enlarged LEs swollen with undegraded cargo (Fig. 1A,B). Actin clearing and ciliogenesis were blocked in CQ-treated cells as in Vps41 KO cells (Fig. 1C-G), suggesting that those phenotypes were indeed due to disruption of lysosomal fusion, rather than a separate non-lysosomal function of Vps41. In both Vps41 KO and CQ-treated cells, actin-clearing defects were accompanied by defects in exclusion of non-ciliary membrane proteins (Fig. 2A) and appeared to be upstream of ciliation defects (Fig. 2B-D), emphasizing the importance of the apical actin cortical clearing step for ciliogenesis in polarized epithelia.

These results align with some previous literature linking lysosomal fusion to ciliogenesis. For example, two inositol-5-phosphatases of which mutations cause cilia defects in patients (INPP5E, involved in Joubert Syndrome, and OCRL, involved in Lowe Syndrome) are important for autophagosome-lysosome fusion (De Leo et al., 2016; Hasegawa et al., 2016; Yamamoto and Mizushima, 2021). Depletion of VAMP7, a SNARE protein involved in lysosomal fusion, was also shown to disrupt ciliation in MDCK cells, although depletion of the lysosomal enzyme α-gal A did not affect ciliation (Szalinski et al., 2014), which may suggest that it is specifically the fusion step of lysosomal trafficking, rather than lysosomal function overall, that is important to ciliogenesis.

While these reports, including the experiments shown here, demonstrate the requirement for lysosomal fusion during ciliogenesis, what remains unclear is the mechanism by which lysosomal fusion affects cilia formation. Recent literature suggested a potential reason why lysosomal fusion might be required for ciliogenesis: several studies have shown that certain ciliogenesis inhibitors, including OFD1 (Tang et al., 2013), MYH9 (Yamamoto et al., 2021), and CP110 (Liu et al., 2021), are degraded by selective autophagy to promote ciliogenesis. Since autophagy involves HOPS-dependent autophagosome-lysosome fusion (Jiang et al., 2014), we hypothesized that the ciliogenesis defects of Vps41 KO MDCK cells might be due to a failure to degrade these ciliogenesis inhibitors. However, although autophagy was impaired in Vps41 KO cells (Fig. S1A,B), these cells did not show excess accumulation of MYH9 (Fig. S1C-E), OFD1, or CP110 (Fig. S1D,E). We also did not observe MYH9 accumulation upon treatment with autophagy inhibitor BafA (Fig. S1C), suggesting that MYH9 was not an autophagy cargo in polarized MDCK cells. The difference between these results and those of previous studies (Liu et al., 2021; Tang et al., 2013; Yamamoto et al., 2021) may reflect differences between starvation-induced and starvation-independent ciliogenesis. Serum starvation is commonly used to induce ciliogenesis in cultured cells, especially in non-polarized cells that use an intracellular ciliogenesis pathway. However, serum starvation also induces autophagy, and it was under starvation conditions that autophagy of MYH9, OFD1, and CP110 was shown to be important for ciliogenesis (Liu et al., 2021; Pierce and Nachury, 2013; Tang et al., 2013; Yamamoto et al., 2021). In contrast, our study examined ciliogenesis in polarized MDCK monolayers that use an extracellular ciliogenesis pathway and do not require starvation. Thus, selective autophagy of ciliogenesis inhibitors may be required for starvation-induced ciliogenesis in some cell types, but not for ciliogenesis in polarized epithelial cells.

### Inhibition of lysosomal fusion disrupts recruitment of Rab19 to the site of ciliogenesis

In searching for alternate explanations for the actin-clearing and ciliogenesis defects of Vps41 KO MDCK, we investigated Rab19, since previous work had shown that Rab19 drives actin-clearing during ciliogenesis and that the HOPS complex is a Rab19 effector (Jewett et al., 2021). As previously reported (Jewett et al., 2021), in WT MDCK monolayers, Rab19 localized to the actin cortical clearing around the basal body and only occasionally localized to LE/Ls (Fig. 3A-D). Rab19 was also observed in the Golgi apparatus (Fig. S2A,B), suggesting that the role of Rab19 in ciliogenesis might involve transporting some ciliogenesis factors from the Golgi to the basal body. In Vps41 KO or CQ-treated cells, however, Rab19 accumulated on membranes of enlarged LEs and its recruitment to the basal body and Golgi was reduced (Fig. 3A-D and Fig. S2A,B).

In support of a hypothesis that depletion of Rab19 from the site of ciliogenesis contributes to the actin-clearing and ciliogenesis defects of Vps41 KO and CQ-treated cells, we found that GFP-Rab19 overexpression rescued those defects (Fig. 3E,F). Thus, retention of Rab19 on LEs in Vps41 KO and CQ-treated cells reduces the fraction of cellular Rab19 that can be recruited to the basal body, thereby disrupting actin-clearing and ciliogenesis (Fig. 6). Interestingly, GFP-Rab19 overexpression only partially rescues ciliation in Vps41 KO cells, while fully rescuing the phenotype in CQ-treated cells (Fig. 3E,F), suggesting that in Vps41 KO cells a reduction of Rab19 at the basal body contributes to but is not the sole reason for ciliation defects. It may be that HOPS complex depletion also affects the localization of other, as yet unidentified, ciliogenesis regulators.

### Rab19 function at LEs

Besides revealing a mechanism by which actin-clearing and ciliogenesis was disrupted in HOPS-depleted cells, the observation that Rab19 accumulates on LEs when lysosomal fusion is inhibited (Fig. 3C,D) also suggested that Rab19 may have a previously uncharacterized function at the LE. Rab19 KO did not affect LE/L size, but expression of the Rab19-CA mutant exacerbated the enlargement of LEs upon CQ treatment (Fig. S2C,D). One potential explanation is that Rab19 may mediate the transport of certain cargoes to LEs, which under normal conditions constitute only a minor fraction of total LE cargo volume, but when downstream lysosomal degradation is blocked then an increase in delivery of these Rab19 cargoes results in increased swelling of the LEs. Further studies will be needed to identify these Rab19-dependent LE cargoes. Rab19 was previously reported to localize to the Golgi and trans-Golgi network (TGN) in *Drosophila* (Sinka et al., 2008), and we confirmed that Rab19 is also found in the Golgi in MDCK cells (Fig. S2A,B), so it may be that Rab19 mediates a TGN-to-LE/L transport route.

The HOPS complex is involved in TGN-to-LE/L transport of certain lysosomal membrane proteins via the AP-3 pathway (Balderhaar and Ungermann, 2013; Schoppe et al., 2020), and proteomics for Rab19 binding partners suggest Rab19 may also interact with the AP-3 complex (Jewett et al., 2021). This raises an interesting possibility, which remains to be tested by future studies, that Rab19-HOPS interaction could be involved in AP-3-dependent TGN-to-LE/L transport. We furthermore found that Rab19 interacts with V-ATPase (Fig. S2F). Since V-ATPase can act as a pH sensor to recruit trafficking factors to endo/lysosomes in an acidification-dependent manner (Hurtado-Lorenzo et al., 2006), this interaction could potentially be involved in targeting Rab19 to the LE or in releasing Rab19 from the lysosome following lysosomal fusion.

### Differential roles of LE/L-associated HOPS complex and EE-associated CORVET complex in ciliogenesis

HOPS is a large protein complex that was shown to bind to other Rab GTPases, specifically to Rab7 which interacts with the HOPS-specific subunits, Vps41 and Vps39. We investigated the interaction between Rab19 and HOPS, aiming to determine which of the HOPS subunits was responsible for binding Rab19. Surprisingly, we found that this interaction was independent of Vps41 and Vps39 and appeared instead to be mediated by the core complex (Vps11, Vps16, Vps18, and Vps33A) that is shared between LE/L-associated HOPS and EE-associated CORVET (Fig. 4 and Fig. S3). Importantly, this revealed that Rab19 may interact with CORVET on EEs as well as with HOPS on LE/Ls. This led us to probe the respective roles of EEs and LE/Ls in ciliogenesis.

The original work that identified HOPS as being required for ciliogenesis (Jewett et al., 2021) left open the question of whether HOPS is actually localized to the basal body to participate directly in ciliogenesis. To address the question of HOPS localization, we first used immunostaining for the HOPS/CORVET core subunit Vps11 and found that Vps11-positive vesicles were frequently present at the basal body (Fig. 5A,B,D). To determine whether the Vps11 at the basal body was part of the LE/L-associated HOPS complex or part of the EE-associated CORVET complex, we used LysoTracker as a marker for LE/Ls and EEA1 immunostaining as a marker for EEs. EEA1-positive EEs were often observed at the basal body and actin clearing, while LysoTracker-positive LE/Ls were not (Fig. 5B,D,F,G). Thus, it appears that the CORVET rather than HOPS is present at the site of ciliogenesis. A role for EEs at the basal body is consistent with previous studies which observed EEA1-positive or Rab5-positive EEs at the base of cilia in various cell types including RPE1 cells, fibroblast-like synoviocytes, astrocytes, and *C. Elegans* sensory neurons (Blacque et al., 2018; Hu et al., 2007; Leitch et al., 2014; Moser et al., 2009; Rattner et al., 2010). This has been interpreted as representing a zone of endocytosis around the base of the cilium, which in these cell types is embedded in a membrane invagination termed the “ciliary pocket” (Molla-Herman et al., 2010; Moser et al., 2009; Rattner et al., 2010). Cilia of polarized epithelial cells, such as the MDCK cells in this study, have been described as protruding directly from the plasma membrane without a ciliary pocket (Ghossoub et al., 2011; Molla-Herman et al., 2010). However, our observations may indicate that these cell types nonetheless have a comparable endocytosis domain around the base of the cilium. It remains unclear whether these EEs are required for ciliogenesis, with some studies finding that ciliation is not impaired by mutations of the EE regulator Rab5 or other early endocytic machinery (Blacque et al., 2018). There is, however, evidence that endocytic trafficking at the base of cilia regulates ciliary membrane receptor localization and signaling functions (Blacque et al., 2018; Clement et al., 2013); we would speculate that such may be the role of the CORVET-containing EEs at the basal body in MDCK cells.

### Conclusions and biomedical implications

Collectively, our observations favor a model in which the HOPS complex is not directly involved in ciliogenesis at the basal body (Fig. 6). Instead, HOPS-mediated lysosomal fusion occurring elsewhere in the cell maintains the flow of endolysosomal trafficking, which is needed to balance Rab19 activity between the separate functions of Rab19 in ciliogenesis and in LE cargo transport. Under normal conditions, a fraction of Rab19 is transiently recruited to LEs and then cycles off of these membranes after lysosomal fusion, keeping a large pool of Rab19 available to drive actin-clearing and ciliogenesis at the basal body (Fig. 6A). When lysosomal fusion is impaired, as in Vps41 KO or CQ-treated cells, then Rab19 is abnormally retained on LEs which fail to fuse to lysosomes; this depletes the pool of Rab19 available to be recruited to the basal body, and thereby disrupts Rab19-mediated actin-clearing and ciliogenesis (Fig. 6B).

These results highlight a relatively unexplored way that defects in lysosomal trafficking can disrupt other cellular processes such as ciliogenesis: besides blocking the degradation of lysosomal cargoes, such defects also dysregulate the activity of cellular machinery that is normally shared between endolysosomal functions and other functions. While we focused here on the mislocalization of Rab19 and its effect on ciliogenesis, it would be interesting for future studies to investigate what other factors are abnormally retained on LE/Ls when their fusion is impaired and what other cellular functions may be dysregulated thereby.

This mechanism may be relevant to certain ciliopathies and other medical conditions involving defects in lysosomal fusion machinery. As mentioned above, these include Joubert Syndrome caused by INPP5E mutation and Lowe Syndrome caused by OCRL mutation (De Leo et al., 2016; Hasegawa et al., 2016); both of these genetic disorders manifest with ciliary defects and the causative genes are important for lysosomal fusion. For mutations in Vps41 and other HOPS subunits, the main disease manifestations are neurological disorders often involving dystonia (Monfrini et al., 2021a; Monfrini et al., 2021b; Sanderson et al., 2021; Steel et al., 2020; van der Welle et al., 2021). These HOPS mutations cause lysosomal abnormalities in fibroblasts (Monfrini et al., 2021a; Steel et al., 2020; van der Welle et al., 2021), but the mechanism linking those lysosomal defects to the dystonia phenotype is unclear. Investigating whether Rab19 is mislocalized and whether cilia are impaired by these HOPS mutations could inform potential treatments for HOPS-associated neurological disorders. Besides genetic diseases, other conditions can also impair lysosomal fusion and thus might impact cilia by the mechanism outlined here. For example, CQ and the related drug hydroxychloroquine (HCQ), which similarly inhibits autophagosome-lysosome fusion (Mondal et al., 2022), are used to treat malaria and certain autoimmune diseases; these drugs can cause adverse effects including neuropsychiatric events, cardiotoxicity, and retinopathy (Doyno et al., 2021; Schrezenmeier and Dörner, 2020). In light of our results showing that CQ blocks ciliogenesis, it will be important to assess whether cilia are altered in patients treated with CQ or HCQ, since cilia dysfunction could contribute to adverse effects of these drugs. Thus, the indirect role of lysosomal fusion in regulating Rab19 activity for ciliogenesis is potentially relevant to a variety of diseases ranging from genetic disorders to malaria.

## MATERIALS AND METHODS

### Cell lines and cell culture

All of the experiments, except where otherwise noted, were performed in polarized epithelial Madin Darby canine kidney cells (MDCK.2; #CRL-2936, ATCC, Manassas, VA, USA). The human embryonic kidney epithelial-like HEK293T cell line (#CRL-3216, ATCC) was used for the siRNA and Vps8-GFP experiments, and for lentiviral vector production to generate stable MDCK cell lines by lentiviral transduction. Cells were maintained at 37°C with 5% CO_2_ in complete media consisting of DMEM (#10-017-CV, Corning, Corning, NY, USA) with 10% FBS (#PS-300, Phoenix Scientific, San Marcos, CA, USA) and 1x penicillin-streptomycin (#30-002-CL, Corning).

The generation of a parental MDCK line stably expressing Tet-inducible Cas9, and the generation of the Vps41 and Rab19 CRISPR KO lines on that background, was previously described (Jewett et al., 2021). The control cell line referred to as “WT MDCK” throughout the manuscript is the Tet-inducible Cas9 parental line, to provide an appropriately matched control for the CRISPR knockout lines. The cell lines labeled as Vps41 KO#1 and Vps41 KO#2 are two clonal lines of Vps41 KO MDCK. The MDCK cell lines stably overexpressing GFP-Rab19 on the WT or Rab19 KO background were previously described (Jewett et al., 2021). The MDCK cell lines stably overexpressing GFP-Rab19 on the Vps41 KO background, and GFP-MYH9 on the WT or Vps41 KO background, were generated by the same approach, using lentiviral transduction and puromycin selection.

For experiments with CQ, the cells were treated with 10 µM chloroquine diphosphate salt (#200-055-2, Sigma, St. Louis, MO, USA) in the media, beginning approximately 0.5-2 hours after seeding the cells for the experiment, and refreshed daily. The experiments with BafA in Fig. S1 used 100 nM bafilomycin A1 (#S1413, Selleck Chemicals, Houston, TX, USA) in the media for the final 16 hrs. For the serum-starved conditions in Fig. S1A, complete media was replaced with serum-free DMEM for 48 hours.

### Plasmids

The pLVX-GFP-Rab19 lentiviral transfer plasmid used to establish the GFP-Rab19 stable cell lines was previously described (Jewett et al., 2021). A pLVX-GFP-MYH9 transfer plasmid was cloned by ligating a NdeI/SalI-digested GFP-MYH9 cassette from CMV-GFP-NMHC II-A (gift from Robert Adelstein, Addgene plasmid #11347) into NdeI/XhoI-digested pLVX-Puro vector. For lentiviral vector production, the transfer plasmid was co-transfected with the packaging plasmid pΔ8.9 and pseudotyping plasmid pVSV-G into HEK293T cells.

The pGEX-KG-Rab19 plasmid, encoding GST-tagged Rab19 for expression in *E. coli* for recombinant protein production, was previously described (Jewett et al., 2021).

### Antibodies and dyes

**Table 1:** Antibodies and fluorescence dyes used for immunofluorescence and western blotting.

| Primary antibodies | Source | Product number | Application |
| --- | --- | --- | --- |
| γ-Tubulin | Sigma-Aldrich | #T5326 | IF |
| Acetyl-α-Tubulin | Cell Signaling Technology (Danvers, MA, USA) | #5335 | IF |
| Arl13b | Antibodies Inc (Davis, CA, USA) | #73-287 | IF |
| LC3B | Novus Bio (Centennial, CO, USA) | #NB600-1384 | WB and IF |
| MYH9 | Sigma-Aldrich | #M8064 | WB |
| OFD1 | Abcam (Waltham, MA, USA) | #ab222837 | WB |
| CP110 | ProteinTech (Rosemont, IL, USA) | #12780-1-AP | WB |
| GM130 | BD Biosciences (San Jose, CA, USA) | #610822 | IF |
| Vps11 | Santa Cruz Biotechnology (Dallas, TX, USA) | #sc-515094 | WB and IF |
| Vps41 | Santa Cruz Biotechnology | #sc-377271 | WB |
| ATP6V1A | Novus Bio | #NBP2-55148 | WB |
| β-tubulin | LI-COR (Lincoln, NE, USA) | #926-42211 | WB |
| α-Tubulin | Sigma-Aldrich | #T6074 | WB |
| α-Tubulin | Santa Cruz Biotechnology | #sc-23948 | WB |
| GFP | Thermo Fisher Scientific (Waltham, MA, USA) | #A11122 | WB |
| Secondary antibodies | Source | Product number | Application |

| AF488 anti-mouse | Jackson ImmunoResearch (West Grove, PA, USA) | #715-545-150 | IF |
| --- | --- | --- | --- |
| AF594 anti-mouse | Jackson ImmunoResearch | #715-585-150 | IF |
| AF488 anti-rabbit | Jackson ImmunoResearch | #711-545-152 | IF |
| AF594 anti-rabbit | Jackson ImmunoResearch | #711-585-152 | IF |
| AF647 anti-rabbit | Thermo Fisher Scientific | #A-21245 | IF |
| AF488 anti-Mouse IgG2a | Thermo Fisher Scientific | #A-21131 | IF |
| AF568 anti-Mouse IgG1 | Thermo Fisher Scientific | #A-21124 | IF |
| IRDye 680RD anti-mouse | LI-COR | #926-68072 | WB |
| IRDye 800CW anti-rabbit | LI-COR | #926-32213 | WB |
| Dyes | Source | Product number | Application |
| LysoTracker Red DND-99 | Thermo Fisher Scientific | #L7528 | IF |
| Hoechst | AnaSpec (Fremont, CA, USA) | #AS-83218 | IF |
| Phalloidin AF647 | Thermo Fisher Scientific | #A22287 | IF |

#### Western blotting

For western blotting of cell lysates, MDCK samples cultured at high confluency were washed with phosphate buffered saline (PBS) and lysed in a buffer of Tris buffered saline (TBS) with 1% Triton X-100 and 1 mM EDTA for 30 min on ice, then centrifuged for 20 min at 15,000✕g at 4°C. The supernatants of the cell lysates were normalized for total protein concentration according to Bradford assay (#5000006, Bio-Rad, Hercules, CA, USA), mixed with SDS loading dye, and boiled for 5-10 min at 95°C. The samples were then run on SDS-PAGE, transferred to PVDF membranes, blocked for at least 20 min in a buffer consisting of Intercept (TBS) Blocking Buffer (#927-60001, LI-COR) diluted 1:3 in TBS with 0.05% Tween-20 (TBST), probed with the primary antibodies diluted in blocking buffer for at least 2 hours at room temp or overnight at 4°C, washed 4x ≥ 5 min with TBST, probed with the secondary antibodies diluted in blocking buffer for 1-2 hours at room temp, washed 4x ≥ 5 min with TBST and once with TBS, and then imaged on an Odyssey DLx imaging system (LI-COR).

Densitometry of the blot images was performed in Image Studio Lite Version 5.2.5 (LI-COR) using median background subtraction. Detected levels of the target protein were normalized according to a tubulin loading control, then normalized to the level in detected in the WT MDCK control in the given experiment.

#### GST-Rab19 pull-down assays

GST-tagged Rab19 protein was recombinantly produced in BL21 Codon Plus *E. coli*, purified using glutathione agarose beads and eluted with free glutathione as previously described (Jewett et al., 2021), and used for glutathione bead pull-down assays with cell lysates, as previously described (Jewett et al., 2021; Willenborg et al., 2011). The bead eluates and cell lysates were then analyzed by SDS-PAGE and western blotting as described above.

#### siRNA knockdowns of HOPS subunits

The siRNAs (listed below) were purchased from Qiagen (Germantown, MD, USA) and were transfected into HEK293T cells using Lipofectamine RNAiMAX transfection reagent (#13778075, Thermo Fisher Scientific), and samples were harvested 2 days post transfection. For RT-qPCR validation of the siRNA knockdowns, RNA was isolated using TRIzol reagent (#15596026, Thermo Fisher Scientific), cDNA was synthesized using the SuperScript IV First-Strand Synthesis System (#18091050, Thermo Fisher Scientific), and qPCR was performed using iTaq Universal SYBR Green Supermix (#1725121, Bio-Rad), with primers (listed below; purchased from Integrated DNA Technologies, Coralville, IA, USA) designed to span an exon-exon junction to prevent amplification from genomic DNA, and analyzed using a StepOnePlus Real-Time PCR machine (#4376600, Thermo Fisher Scientific). GAPDH was analyzed as a housekeeping gene control.

**Table 2:**
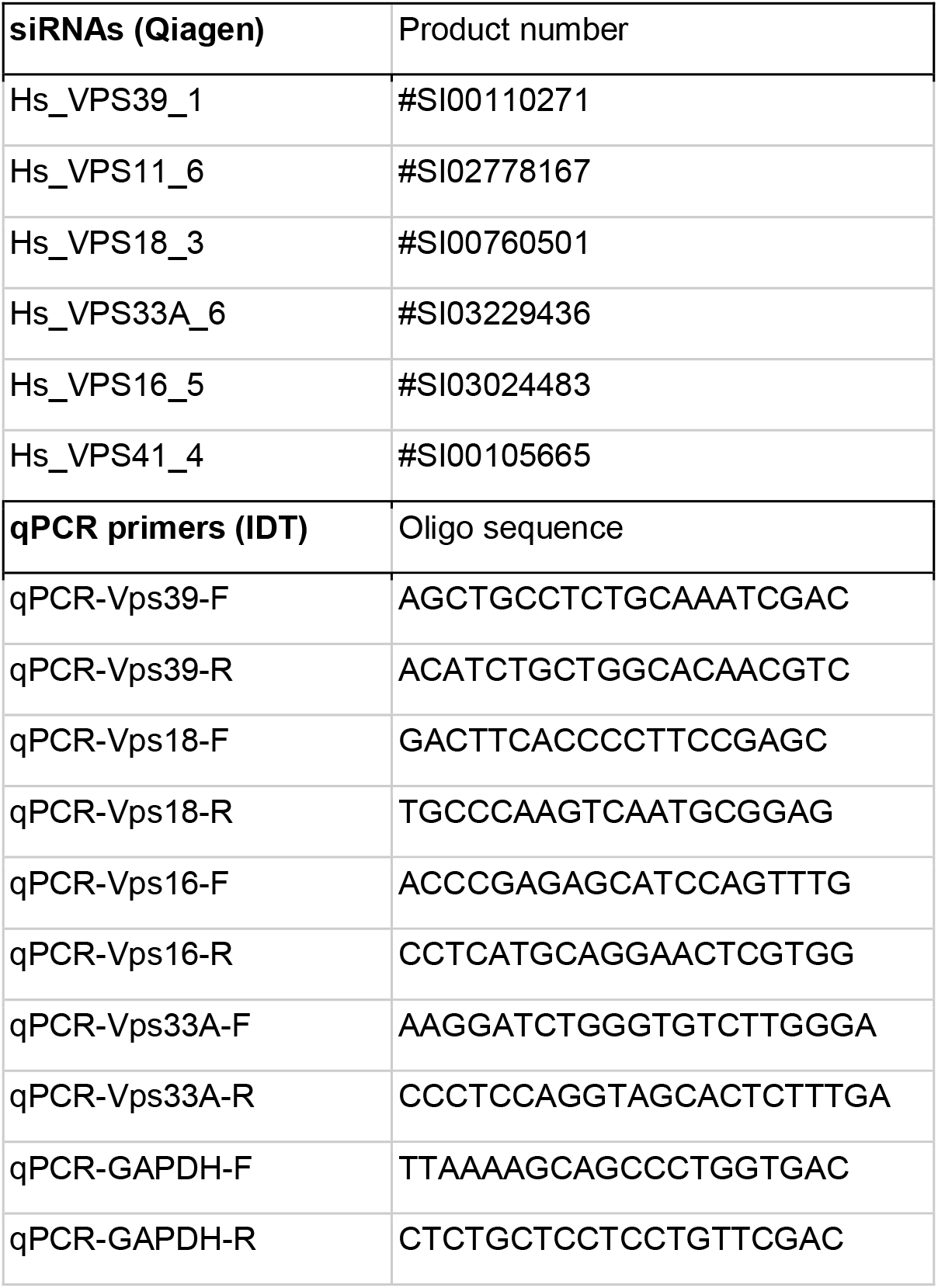
siRNAs and qPCR primers

#### GFP-Rab19 pull-down assay

MDCK cells expressing GFP-Rab19 on the Rab19 KO background were lysed and incubated with GST-tagged anti-GFP nanobody (recombinantly produced from pGEX6P1-GFP-Nanobody, Addgene Plasmid #61838) or with free GST for negative control, followed by GTPγS or GDP loading and GST pulldown as previously described (Jewett et al., 2021), and the IP eluates were analyzed by western blotting for ATP6V1A.

#### Immunofluorescence experiments

##### Culturing of MDCK monolayers on transwell filters

µm pore PET membrane transwell filter inserts (12-well filters, #665640, Greiner Bio-One, Monroe, NC, USA; or 24-well filters, #662640, Greiner Bio-One) in wells of tissue-culture well plates were pre-coated with rat-tail collagen and cured under UV light for ≥ 30 min prior to use. MDCK cells were plated on the filters at a highly confluent density of approximately 2.2✕10^5^ cells/cm^2^ (2.5✕10^5^ cells per 12-well filter, or 7.4✕10^4^ cells per 24-well filter). The cells were cultured on the filters, in complete media which was changed daily, for 3 days prior to fixation for experiments assessing ciliation, actin cortical clearing, and basal body localization phenotypes; or 2 days for some of the other experiments such as those assessing LE/L phenotypes. For experiments with LysoTracker, the live cells were stained with LysoTracker Red DND-99 (#L7528, Thermo Fisher Scientific) at 500 nM (1:2000 dilution) in complete media for the final 30 min prior to fixation.

##### Immunostaining and confocal fluorescence microscopy

The following immunostaining protocol was performed at room temp. Cells on transwell filters in well plates were fixed with 4% paraformaldehyde (#15710, Electron Microscopy Sciences, Hatfield, PA, USA) in PBS for 20 min, quenched 2✕ 5 min with 0.1 M glycine in PBS, and rinsed with PBS. The filters were then cut out of the transwell inserts and placed on parafilm in a humidified dish for the following steps. Filters were rinsed with PBS with 0.1% Triton X-100 (PBST), blocked with 10% normal donkey serum (#017-000-121, Jackson ImmunoResearch) in PBST for ≥ 1 hr, and stained overnight with primary antibodies (listed above) diluted in blocking buffer. The next day, the filters were washed 5✕ ≥ 5 min with PBST, stained with Hoechst and dye-labeled secondary antibodies (listed above) and/or Phalloidin for 2 hrs, washed 5✕ ≥ 5 min with PBST and once with PBS, then mounted on slides (#TNR WHT90, Tanner Scientific, Sarasota, FL, USA) with Vectashield (#H-1000, Vector Laboratories, Newark, CA), covered with coverslips (#48366-067, VWR, Radnor, PA, USA) and sealed with clear nail polish. Confocal fluorescence microscopy images were acquired either on a Nikon A1R confocal microscope (Nikon, Tokyo, Japan) using a 60x oil objective and NIS-Elements AR software (Nikon), or on a Leica SP8 confocal microscope (Leica Microsystems, Wetzlar, Germany) using a 63x oil objective and LAS X software (Leica).

##### Quantitative image analysis

###### LysoTracker compartments (LE/L) size analysis

Z-stack images of MDCK monolayers (grown for 2 days on filters, and stained with LysoTracker) were acquired on the Nikon A1R confocal microscope, with 0.21 µm pixel size and z-step. Analysis of LE/L size was performed in FIJI (Schindelin et al., 2012), as follows: 50 µm ✕ 50 µm ROIs were cropped from the LysoTracker Red channel image stack. The image stack was smoothed, a maximum intensity projection was taken, and rolling-ball background subtraction was applied with a radius of 20 pixels. The image was then binarized using the Yen thresholding algorithm. Particle analysis was applied to quantify the size of LysoTracker compartments, analyzing particles with size ≥ 3 pixels and circularity 0.50-1 and excluding those on edges. Between 6-8 technical replicates (ROIs) were averaged from each biological replicate for each condition. Outlier biological replicates were excluded (ROUT, Q=2%).

###### Measuring apical cortical actin clearing

Z-stack images of MDCK monolayers (grown for 3 days on filters, and stained with anti-γ-tubulin antibody and Phalloidin) were acquired on the Nikon A1R confocal microscope, with 0.28 µm pixel size and z-step. Analysis of apical actin cortical clearing was performed in FIJI, as follows: The z-stack was resliced to obtain the side view. Individual basal bodies (marked by γ-tubulin) docked at the apical actin cortex (marked by Phalloidin) were selected for analysis. An 8 µm wide ✕ 5 µm high ROI was drawn centered around the selected basal body. The Plot Profile function was used to measure the vertically averaged pixel intensities in the γ-tubulin and Phalloidin channels along the horizontal distance through the ROI. Each intensity profile was normalized to have mean=1, by dividing each intensity value by the mean of that entire profile. To determine the fraction of cortical actin remaining over the basal body, the mean of the values in the actin intensity profile within 0.3 µm horizontal distance from the center of the ROI was divided by the mean of the values in the actin intensity profile between 2-4 µm from the center of the ROI. Between 10-15 technical replicates (individual basal bodies) were averaged from each biological replicate for each condition.

###### Measuring ciliation in MDCK cells

Z-stack images of MDCK monolayers (grown for 3 days on filters, and stained with anti-Arl13b antibody and Hoechst) were acquired on the Nikon A1R confocal microscope, with 0.28 µm pixel size and z-step. Analysis of the percentage of cells exhibiting an Arl13b-positive primary cilium was performed in FIJI, as follows: To count cilia, a maximum intensity projection of the Arl13b channel stack was taken, and rolling-ball background subtraction was applied with a radius of 100 pixels. The image was binarized using the Shanbhag thresholding algorithm, and particle analysis was applied to quantify the number of Arl13b puncta, analyzing particles with size ≥ 2 pixels. To count cells, a maximum intensity projection of the Hoechst channel stack was taken, the image was smoothed, and rolling-ball background subtraction was applied with a radius of 100 pixels. The image was binarized using the MinError(I) thresholding algorithm, and watershed separation was used to distinguish overlapping nuclei. Particle analysis was applied to quantify the number of nuclei, analyzing particles with size ≥ 150 pixels. The number of cilia was then divided by the number of nuclei to calculate the percent ciliation. 3-4 technical replicates (142 µm ✕ 142 µm fields of view) were averaged from each biological replicate for each condition.

###### Quantifying Rab19 enrichment at the basal body

Z-stack images of GFP-Rab19-expressing MDCK monolayers (grown for 3 days on filters, and stained with anti-ɣ-tubulin antibody and Phalloidin) were acquired on the Leica SP8 confocal microscope, with 0.12 µm pixel size and z-step, spanning from above the apical cortex to a mid-basolateral plane of the monolayer. Analysis of the enrichment of Rab19 vesicles at the basal body was performed in FIJI, as follows: The z-stack was resliced to obtain the side view. Rolling ball background subtraction was applied with a radius of 75 pixels. Side view slices showing apically docked basal bodies were selected for analysis, and GFP-Rab19 signal in the image slice was binarized using the Yen thresholding algorithm. The binarized result was visually inspected to check that it was a reasonable representation of the Rab19 compartments in the original image. Any cells for which almost all pixels were thresholded as either uniformly positive or uniformly negative (which could occur when GFP-Rab19 expression in the cell of interest was far brighter or far dimmer than in the other cells in the slice) were excluded from analysis. A “basal body” ROI of 3 µm ✕ 2 µm was then drawn with its upper edge centered at the apical end of the basal body, and a “cytosol” ROI was drawn freehand to enclose all of the cytosol of the selected cell that was visible in the image slice, bounded by the Phalloidin-stained actin cortex and excluding the nucleus. The density of Rab19-positive pixels (number of positive pixels divided by total pixel area of the ROI) in the basal body ROI was divided by the density of Rab19-positive pixels in the cytosol ROI to obtain a measure of Rab19 enrichment at the basal body. 10 technical replicates (individual cells) were averaged from each biological replicate for each condition.

###### Analyzing Rab19 colocalization with LysoTracker

Z-stack images of GFP-Rab19-expressing MDCK monolayers (grown for 2 days on filters, and stained with LysoTracker) were acquired on the Leica SP8 confocal microscope, with 0.12 µm pixel size. Background subtraction and binarization of the GFP-Rab19 and LysoTracker image channels was performed in FIJI, as follows: Rolling ball background subtraction was applied with a radius of 100 pixels. 30 µm ✕ 30 µm ROIs were cropped from the image stack, and a single slice at a mid-apical level of the monolayer was selected for analysis. The image slice was binarized using the Yen thresholding algorithm. The fractional overlap (Manders’ Colocalization Coefficients) (Manders et al., 1993) between GFP-Rab19 and LysoTracker was then computed using a custom script written in MATLAB (MathWorks, Natick, MA, USA). 4 technical replicates (ROIs) were averaged from each biological replicate for each condition.

###### Quantifying EEA1 and LysoTracker enrichment at the basal body

Z-stack images of MDCK monolayers (grown for 3 days on filters, and stained with anti-D-tubulin antibody and anti-EEA1 antibody and/or LysoTracker) were acquired on the Leica SP8 confocal microscope, with 0.12 µm pixel size and z-step, spanning from above the apical cortex to a mid-basolateral plane of the monolayer. Analysis of the enrichment of EEA1 and/or LysoTracker vesicles at the basal body was performed by the same method as described above for analysis of Rab19 enrichment at the basal body. 15 technical replicates (individual cells) were averaged from each biological replicate for each condition.

###### Analyzing Vps11 colocalization with EEA1 and with LysoTracker

Z-stack images of MDCK monolayers (grown for 3 days on filters, and stained with Vps11 antibody and either EEA1 antibody or LysoTracker) were acquired on the Leica SP8 confocal microscope, with 0.12 µm pixel size. Background subtraction and binarization, followed by computation of the fractional overlap in a selected z-slice at a mid-apical level of the monolayer, was performed by the same methods as described for Rab19/LysoTracker colocalization above.

###### Statistical analysis

GraphPad Prism 9.5.0 software (GraphPad Software, Boston, MA, USA) was used to generate graphs and to calculate P-values. For all of the quantitative image analysis experiments, each data point on the graphs in the figures represents the mean of the technical replicates (ROIs or individual cells) for one biological replicate (separate experiment), except as noted for the graph in Fig. 2B where the data points represent individual technical replicates. P-values were calculated from the means for the biological replicates. For the analysis of percent ciliation, statistical significance was assessed by mixed-effects analysis with Geisser-Greenhouse correction and Šidák’s multiple comparisons test. For the analysis of basal-body enrichment of EEA1 and LysoTracker, statistical significance was assessed by Kolmogorov-Smirnov test. For the other quantitative image analysis experiments, statistical significance was assessed by Brown-Forsythe and Welch ANOVA test and Dunnett’s T3 multiple comparisons test. For the western blot densitometry results, each data point on the graphs represents the result from one biological replicate (separate set of cell lysates), normalized to the loading control and then to the WT level of the target protein from the same biological replicate, and statistical significance was assessed by one-sample t-test.

## Abbreviations used

HOPS: homotypic fusion and protein sorting;
KO: knockout;
LE: late endosome;
LE/L: late endosome / lysosome,
CORVET: class C core vacuole/endosome tethering;
EE: early endosome;
WT: wild-type;
MDCK: Madin-Darby canine kidney;
CQ: chloroquine;
CA: constitutively active;
V-ATPase: vacuolar-type ATPase;
EEA1: early endosome antigen 1;
IP: immunoprecipitation.

## ACKNOWLEDGEMENTS

We thank Dr. Robert Adelstein for the gift of the CMV-GFP-NMHC II-A plasmid (Addgene #11347). This work was funded by NIDDK grant #2R01DK064380-15 to R.P.

